# Metal Ions and their Effects on Antimicrobial Resistance Development in Wastewater

**DOI:** 10.1101/2023.06.16.545339

**Authors:** Indorica Sutradhar, Prinjali Kalyan, Kelechi Chukwu, Akebe Luther King Abia, Joshua Mbanga, Sabiha Essack, Davidson H. Hamer, Muhammad H. Zaman

## Abstract

Antimicrobial resistance (AMR) is a global health challenge and there is increasing recognition of the role of the environment, particularly wastewater, in the development and spread of AMR. Although trace metals are common contaminants in wastewater, the quantitative effects of trace metals on AMR in wastewater settings remain understudied. We experimentally determined the interactions between common antibiotic residues and metal ions found in wastewater and investigated their effects on the development of antibiotic resistance in *Escherichia coli* over time. These data were then used to expand on a previously developed computational model of antibiotic resistance development in continuous flow settings to incorporate the effects of trace metals acting in combination with multiple antibiotic residues. We found that the common metal ions, copper and iron, interact with both ciprofloxacin and doxycycline at wastewater relevant concentrations. This can significantly affect resistance development due to antibiotic chelation of the metal ions causing a reduction in the antibiotics’ bioactivity. Furthermore, modeling the effect of these interactions in wastewater systems showed the potential for metal ions in wastewater to significantly increase the development of antibiotic resistant *E. coli* populations. These results demonstrate the need to quantitatively understand the effects of trace metal-antibiotic interactions on AMR development in wastewater.

## Introduction

Antimicrobial resistance (AMR) is a pressing global health challenge responsible for about 5 million fatalities in 2019 alone, an annual mortality rate expected to double to 10 million/year by 2050 (1). While global efforts in addressing AMR have largely focused on human health and agriculture sectors, there is increasing evidence that the environment also plays a key role in the development and spread of AMR (1). One of the major reservoirs of AMR in the environment is wastewater, where effluent from hospital, pharmaceutical, domestic, and agricultural sources can contain antibiotic residues and/or bacteria containing antibiotic resistance genes (ARGs) (1). While the effects of these antibiotic residues on the development of AMR in wastewater have begun to be studied and computationally modelled (2), there still exists a gap in modeling the effects of other components of the complex wastewater environment and their effects on AMR development.

One of the most common contaminants in wastewater are trace metal ions, and their presence can have significant environmental and human health implications (3). Industrial and domestic activities can lead to the discharge of metals such as zinc, copper, and iron into wastewater (3-4). For instance, studies in Pakistan and Ghana showed iron concentrations in wastewater ranging from 0.4 mg/L to 4.9 mg/L (5, 6). Furthermore, the presence of these heavy metals in wastewater has been increasing in recent years due to industrial growth including the plating and electroplating industry, batteries, pesticides, the mining industries, the textile industry, as well as the metal smelting, and petrochemicals industries (7). Additionally, these metals do not biodegrade and can pose long term threats to environmental and human health (3, 7). These metals commonly found in wastewater can have a significant impact on the development of AMR in the environment. Studies have shown that trace metals, including copper and zinc, at environmentally relevant and sub-inhibitory concentrations, can promote horizontal gene transfer (HGT) of ARGs between *Escherichia coli* strains (3, 8). Additionally, some trace metals can interact with antibiotic residues forming complexes with changed bioactivity (9, 10). For example, studies have found that ciprofloxacin can bind to ferric ion which reduces ciprofloxacin bioavailability (10). Similarly, iron chelation by doxycycline has been seen to decrease the anti-pseudomonal activity of doxycycline (9). However, the quantitative effects of these trace metals on AMR in wastewater settings have not been adequately studied. There is thus a need to incorporate trace metals into models of AMR development in the environment. Our study attempted to fill this gap by experimentally determining the degree of interaction between common antibiotic residues and metal ions seen in wastewater and probing the effect of these interactions on the development of antibiotic resistance over time. Furthermore, we use these results to expand our previously developed computational model of antibiotic resistance development in continuous flow settings to include the effects of trace metals present alongside antibiotic residues on the development of AMR in wastewater (2).

## Methods

### Metal-Antibiotic Interaction Determination

Wild type *E. coli* MG1655 was cultured for 24 hours at 37°C in LB media supplemented with 2-fold concentration increments of ciprofloxacin, doxycycline, erythromycin, streptomycin, and rifampin in 96-well plates. Various increasing concentrations of copper (II) sulfate pentahydrate (Sigma-Aldrich, St. Louis, US) or iron (III) sulfate hydrate (Sigma-Aldrich, St. Louis, US), chosen for their common occurrence in wastewater and their known interactions with antibiotics, were added to each antibiotic condition (3, 8-10). Each combination of metal and antibiotic was run in biological duplicate. The concentrations of metals surveyed were between 0.5 mg/L to 100 mg/L, a range determined to be relevant to wastewater environments (5, 6). The concentration of metal required to double the MIC of *E. coli* to ciprofloxacin or doxycycline as compared to wild type was then used in the developed model to determine the effect of the metal on the effective concentration of antibiotic in the system.

### Serial Passaging

Wild type *E. coli* MG1655 cultured at 37°C in LB media was grown in 96-well plates with 2-fold increments of ciprofloxacin with the maximum concentration at 4.8 ug/mL. Additionally, the metal of interest (0.5 mg/l or 5 mg/L copper or 5 mg/L or 10 mg/L iron) was added to each well. Each combination of metal and antibiotic was run in biological triplicate (n=3). We selected the bacteria in the well closest to 50% of the inhibitory concentration (IC50) to seed new bacterial cultures on the same dose series of ciprofloxacin at ∼ 24 hours, for 10 days. Bacteria serially passaged in media for the duration of the experiments served as the control groups. This procedure was repeated with 2-fold increments of streptomycin with the maximum concentration at 254 ug/mL and 0.05 mg/L, 0.5 mg/L and 5 mg/L copper.

### Computational Model

The model used in this paper is based on a previously developed and validated model of the growth of antibiotic resistant bacterial populations in continuous flow environments (2). This model was built on prior studies and was extended to incorporate a variety of wastewater relevant inputs which can be broadly classified into bacterial parameters, environmental parameters, and antibiotic parameters. Bacteria specific input factors include the growth rates of antibiotic susceptible and resistant strains and mutation rates in response to subinhibitory concentrations of antibiotic. The antibiotic specific inputs, such as bactericidal activity, allow for the study of the effects of antibiotic pollution on the development of resistance. Additionally, environmental inputs, including physical inflow and outflow rates and antibiotic residue concentrations, allow for the modelling of resistance development in a variety of settings of interest. Ordinary differential equations incorporating these input parameters were used to model an output of resistant bacterial populations over time, thus allowing for the prediction of resistant population development. In this work, we expanded the model to include the effects of two metal ions, copper and iron, on resistance development in response to various antibiotic combinations.

This was done by adjusting the effective concentration of each antibiotic by the degree to which the antibiotic chelated the metal ion concentration that was present (Eq set 1). Each antibiotic is now modeled as an “available antibiotic concentration” (*C*_*1*_ and C_*2*_). These concentrations are modelled with experimentally determined terms for the antibiotic residue concentration in the environment (*E*), the antibiotic clearance rate (*k*_*e*_), and chelation of two metal ions (*M*_*1*_ and *M*_*2*_). The degree of chelation (*k*_*M*_) term is determined for each antibiotic-metal pair by calculating fractional amount of antibiotic that can be bound to the given metal ion from the results of the metal-antibiotic interaction determination described above.

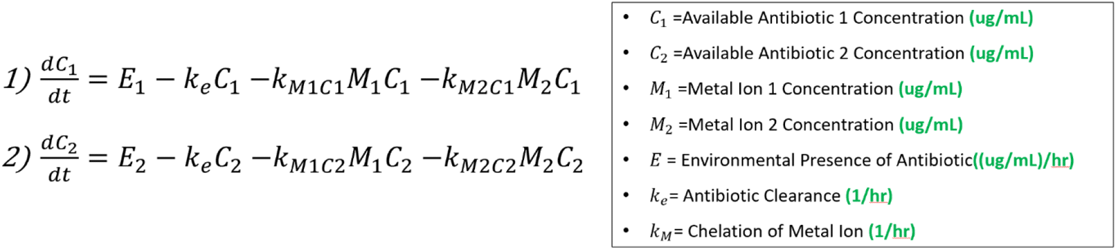

**Eq Set. 1** Model equations for available antibiotic concentrations in the presence of two metal ion concentrations

## Results and Discussion

Initial experiments examining the interactions between various antibiotics and iron and copper solutions demonstrated that only doxycycline and ciprofloxacin showed changes in antibiotic activity against *E. coli* in the presence of the metal ions (Table 1).

**Table 1.** Interactions of various antibiotics with iron and copper solutions

| Antibiotic | Concentration of Iron to Double MIC | Concentration of Copper to Double MIC |
| --- | --- | --- |
| Doxycycline | 5 mg/L | 20 mg/L |
| Ciprofloxacin | 20 mg/L | 50 mg/L |
| Erythromycin | No Interaction | No Interaction |
| Streptomycin | No Interaction | No Interaction |
| Rifampicin | No Interaction | No Interaction |

Due to the comparatively lower change in activity of the ciprofloxacin in response to iron and copper at environmentally relevant concentrations as well as its high relevance in clinical settings in low- and middle-income countries, ciprofloxacin was chosen for longitudinal studies of the effect of these metal ions on resistance development over time. In the first of these experiments, the effect of iron ions alongside ciprofloxacin significantly increased the development of ciprofloxacin resistance in E. coli (Figure 1). This is in agreement with literature findings that iron binding can reduce the bioactivity of ciprofloxacin and evidence that subinhibitory ciprofloxacin concentrations can lead to the development of ciprofloxacin resistance (10, 11).

**Figure 1.**
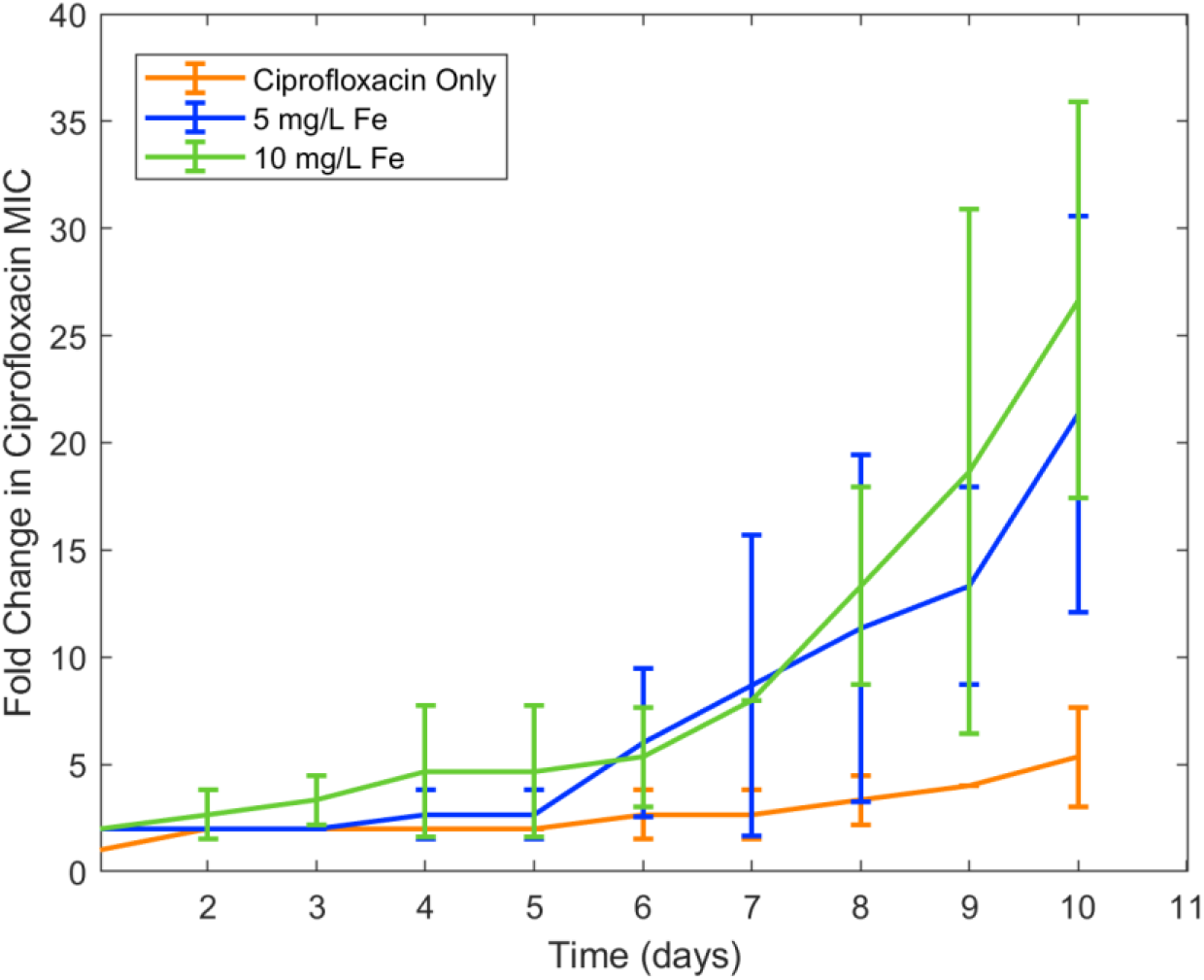
Effect of iron on ciprofloxacin resistance development

Because copper was known to have similar binding to ciprofloxacin as iron, in addition to antimicrobial effect of its own, it was chosen for a second trial of the experiment. Similar results were seen in examining the effect of copper ion alongside ciprofloxacin. While the lower concentration of copper (0.5 mg/L) did not have an effect on resistance development, the higher concentration (5 mg/L) increased ciprofloxacin resistance development in *E. coli* (Figure 2). As with iron, copper has been known to bind with ciprofloxacin resulting in reduced bioactivity, which is likely the cause of this increase in resistance. Unlike iron, copper has antimicrobial activity of its own which may have also contributed to the resistance development; however, at the low concentrations studied, copper activity against *E. coli* was minimal compared to the significant binding between copper and ciprofloxacin that was seen. In order to confirm the effect of chelation compared to the antimicrobial activity of copper, we chose to conduct an experiment with copper and streptomycin, as this combination previously did not show any interaction and no binding between these components have been previously observed. These results showed similar resistance development between the conditions in the presence of copper and those with only streptomycin (Figure 3). Combined with the results of the ciprofloxacin-iron and ciprofloxacin-copper combinations, we can conclude that antibiotic chelation of metal ions can have a significant effect on AMR development over time. Though the effects of metal ions on the transfer of ARGs in wastewater has previously been noted, this mechanism of resistance development in response to metal ions has not previously been observed (3, 8).

**Figure 2.**
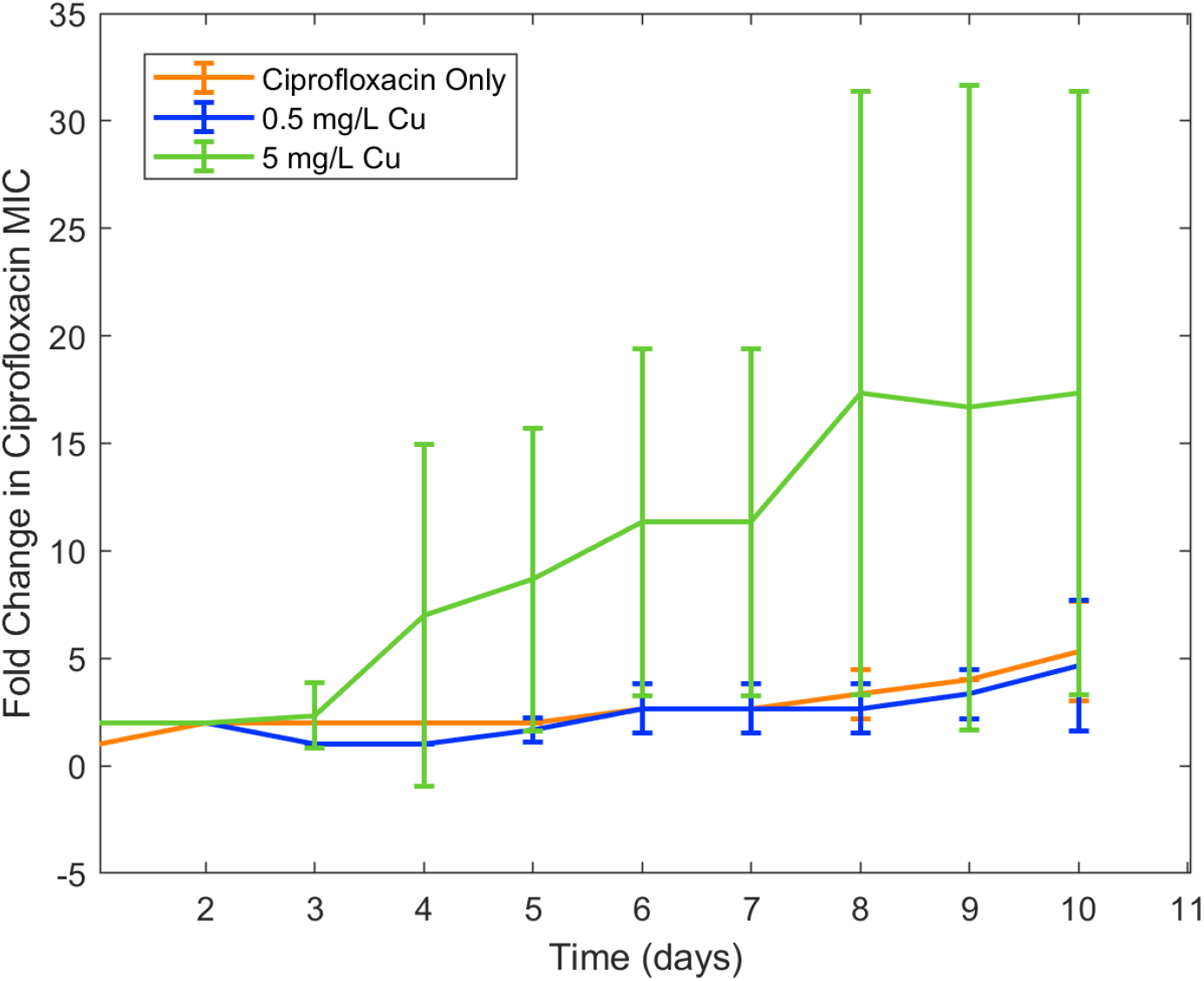
Effect of copper on ciprofloxacin resistance development

**Figure 3.**
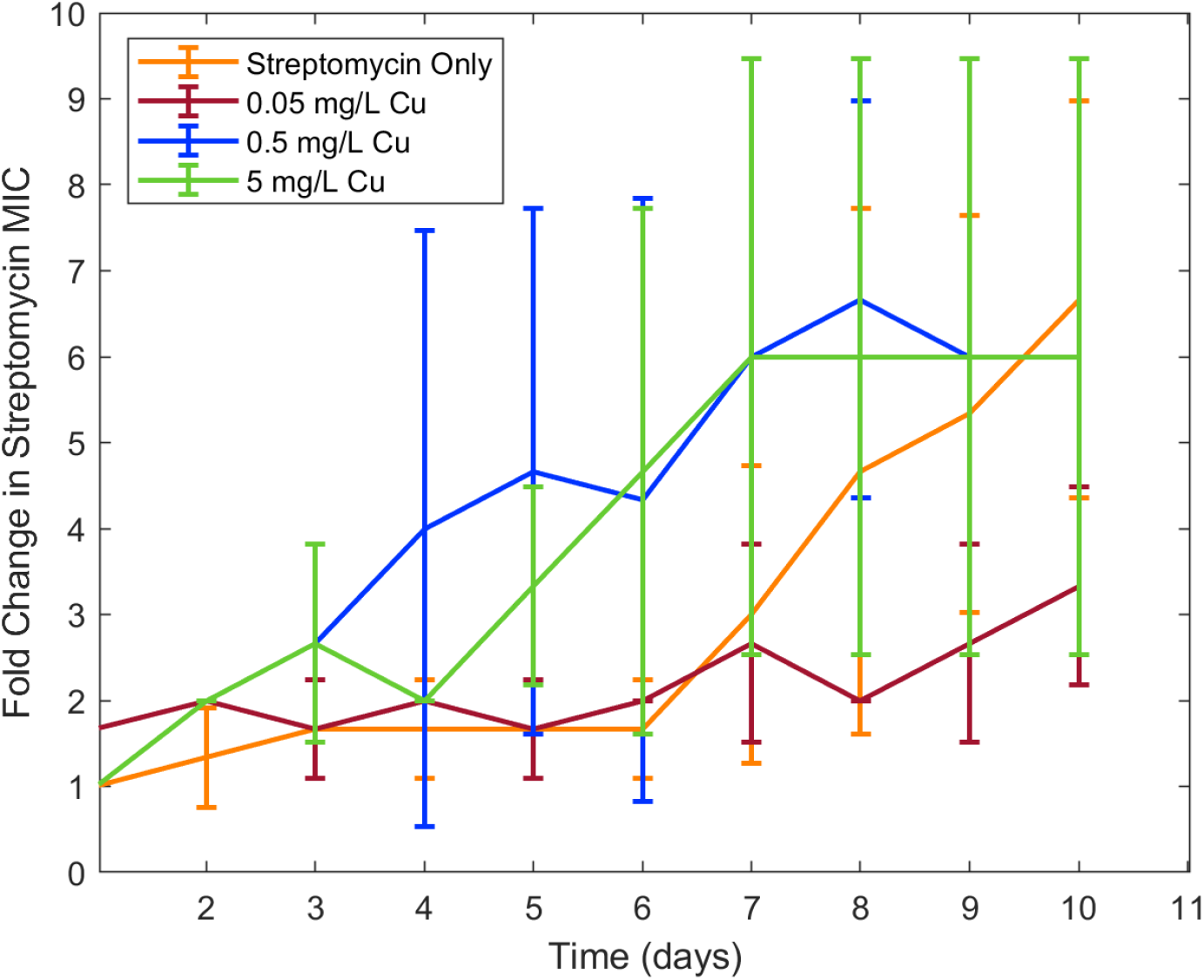
Effect of copper on streptomycin resistance development

In order to probe the potential effects of these metal-antibiotic interactions on the development of AMR in wastewater environments, metal concentrations were added to our previously developed model of antibiotic resistance in wastewater, and the outputs in the presence and absence of metal ion concentrations were compared. Previously, through computational and experimental methods, we developed a model of the antagonistically interacting doxycycline and ciprofloxacin and we found that E. coli populations grown with these antibiotics showed an antibiotic ratio-dependent development of resistance (12). Specifically, conditions with a lower MIC fraction of doxycycline compared to ciprofloxacin showed more resistance development than conditions with an equal ratio or greater relative doxycycline concentration (12). After the addition of iron and copper concentrations to the model, we reexamined the same conditions in the presence and absence of 5 mg/L iron and 5 mg/L copper. Interestingly, in the case of a combination of concentrations of 0.5X MIC ciprofloxacin and 0.5X MIC doxycycline, the addition of iron and copper ions result in the development of a *E. coli* population resistant to both antibiotics that was not seen in the absence of the metal ions (Figure 4). This finding is consistent with both the previously ratio-dependent behavior of the doxycycline-ciprofloxacin combination resistance development and the relatively high doxycycline metal ion chelation observed in this study. Thus, the increased resistance development is due to increased chelation activity of doxycycline, activating the ratio-dependent resistance development behavior noted previously by tipping the antibiotic ratio towards a greater relative ciprofloxacin concentration.

**Figure 4.**
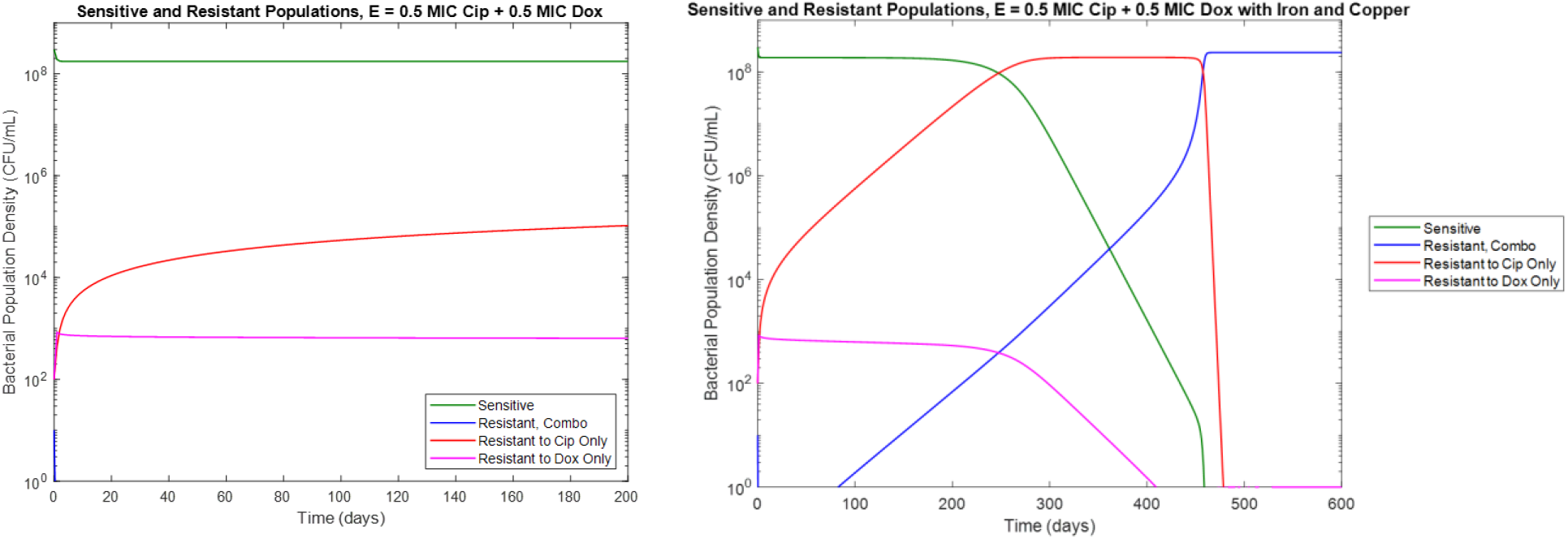
Comparison of resistance development under 0.5X MIC ciprofloxacin and 0.5X MIC doxycycline in the **a.)** absence of iron and copper **b.)** presence of 5 mg/L iron and 5 mg/L copper

## Conclusions

Overall, we found that metal ions, such as iron and copper, commonly found in wastewater, can significantly affect the development of antibiotic resistance for certain types of antibiotics. We observed significant interactions between these metal ions and both ciprofloxacin and doxycycline, leading to changes in the bioactivity of these clinically important antibiotics against *E. coli*. This is in agreement with previous literature on the chelation activity of these antibiotics on these metal ions (9, 10). Furthermore, we observed that this reduction in bioactivity from the presence of iron and copper increased the development of resistance to ciprofloxacin overtime, a novel finding not previously reported in studies of ciprofloxacin-metal interactions. These effects were then incorporated into a computational model of AMR in wastewater. These model simulations showed that metal-antibiotic interactions can significantly increase the development of antibiotic resistance over time in certain conditions. While the effects of metal ions on increased HGT of antibiotic resistance genes in the environment have been recorded, the effect of antibiotic-metal ions binding on resistance acquisition has not previously been observed.

While our studies have demonstrated that metal-antibiotic interaction can significantly affect AMR development in wastewater, we note that our studies have limitations. One limitation was that we only studied a small number of metal-antibiotic combinations and as such cannot conclude that these effects will occur for metal-antibiotic interactions. Future studies investigating a wider array of trace metal ions could further illuminate the full impact of these ions on wastewater AMR. Additionally, we were limited by the infeasibility of experimentally validating our computational findings using continuous flow experimental setups due to the long timeframe and large resource requirements. Further innovation in experimental models of wastewater AMR will be required to fully understand the effects of metal-antibiotic interactions on AMR development.

Despite these limitations, we have demonstrated that metal ions alongside antibiotic residues in wastewater can significantly effect on the emergence of AMR over time. Through both experimental validation and computational modelling, we have observed that interaction between common metal ions and antibiotic residues can result in bioactivity changes leading to increased AMR development. These findings have important implications for understanding the effects of industrial runoff of metal ions in wastewater on AMR and highlight the need to consider the presence of metal ions when assessing the risk of AMR development in wastewater environments. The developed model has potential to be used as a prediction tool for policy makers in both public health and environmental regulation in determining acceptable metal ion and antibiotic residue levels in wastewater systems.

## Acknowledgments

This research was supported in part by the National Institutes of Health training grant at Boston University, T32 EB006359. The content is solely the responsibility of the authors and does not necessarily represent the official views of the National Institutes of Health.

